# BET inhibition rescues ciliogenesis and ameliorates pancreatitis-driven phenotypic changes in mice with Par3 loss

**DOI:** 10.1101/2023.09.14.557654

**Authors:** Mario A. Shields, Anastasia E. Metropulos, Christina Spaulding, Tomonori Hirose, Shigeo Ohno, Thao N.D. Pham, Hidayatullah G. Munshi

**Author notes:** Address correspondence to Mario A. Shields, Ph.D. or Hidayatullah G. Munshi, M.D., Department of Medicine, Northwestern University Feinberg School of Medicine, 303 E. Superior Ave., Lurie 3-220 (M.A. Shields) or Lurie 3-117 (H.G. Munshi), Chicago, IL 60611, USA.; (M.A. Shields); (H.G. Munshi).

## Abstract

The apical-basal polarity of pancreatic acinar cells is essential for maintaining tissue architecture. However, the mechanisms by which polarity proteins regulate acinar pancreas tissue homeostasis are poorly understood. Here, we evaluate the role of Par3 in acinar pancreas injury and homeostasis. While Par3 loss in the mouse pancreas disrupts tight junctions, Par3 loss is dispensable for pancreatogenesis. However, with aging, Par3 loss results in low-grade inflammation, acinar degeneration, and pancreatic lipomatosis. Par3 loss also exacerbates pancreatitis-induced acinar cell loss, resulting in pronounced pancreatic lipomatosis and failure to regenerate. Moreover, Par3 loss in mice harboring mutant Kras causes extensive pancreatic intraepithelial neoplastic (PanIN) lesions and large pancreatic cysts. We also show that Par3 loss restricts injury-induced primary ciliogenesis. Significantly, targeting BET proteins enhances primary ciliogenesis during pancreatitis-induced injury and, in mice with Par3 loss, limits pancreatitis-induced acinar loss and facilitates acinar cell regeneration. Combined, this study demonstrates how Par3 restrains pancreatitis- and Kras-induced changes in the pancreas and identifies a potential role for BET inhibitors to attenuate pancreas injury and facilitate pancreas tissue regeneration.

## INTRODUCTION

Cell polarity plays a crucial role in establishing cellular architecture and function ^1,2^. Proper maintenance of cell polarity is necessary for tissue integrity as cell polarity prevents damage and preserves tissue homeostasis ^3-5^. Several polarity programs control epithelial polarization and morphogenesis ^1,2^. For example, the Par3 complex – comprising partitioning-defective (Par)3, the atypical protein kinase C (aPKC), and Par6 – plays an essential role in controlling the apical domain identity ^3-5^. Accordingly, loss of Par3 in the skin reduces self-renewal, increases differentiation, and causes premature skin aging ^6^. Despite the importance of Par3 in skin homeostasis ^6,7^, the role of Par3 in maintaining other tissue types is poorly understood.

The pancreas comprises the endocrine pancreas and the exocrine pancreas, which produces and secretes digestive enzymes into pancreatic ducts ^8,9^. Animal studies have shown that the exocrine pancreas possesses an intrinsic capacity for regeneration ^8,9^. Induction of acute pancreatitis results in acinar cells losing their differentiation and acquiring a duct-like state in a process called acinar-to-ductal metaplasia (ADM) ^8-10^. During pancreas regeneration, the ADM resolves as the acinar cells re-differentiate ^8-11^. Oncogenic Kras mutation, one of the earliest genetic mutations in human pancreatic cancer ^12,13^, also promotes the development of ADM ^11^. These mutant KRAS-expressing metaplastic ducts can, in turn, progress into precursor lesions known as pancreatic intraepithelial neoplastic (PanIN) lesions ^12,13^. While there are changes in the apical-basal cell polarity during ADM and PanIN formation ^14-16^, the role of polarity proteins in the acinar pancreas homeostasis is yet to be fully understood. Here, we evaluate the role of the polarity protein Par3 in acinar pancreas homeostasis.

## RESULTS

### Increased Par3 expression in human and mouse pancreatitis specimens

Initially, we evaluated the expression of Par3 in chronic pancreatitis specimens using a previously validated Par3 antibody and human tissue microarray slides containing human pancreatitis sections. Compared to mild acute pancreatitis, we found increased expression of Par3 in the chronic pancreatitis specimens (**Fig. 1A**). Furthermore, we found increased expression of Par3 in ADM lesions present in the human pancreatitis specimens compared to the surrounding pancreas (**Fig. 1B**). We also assessed Par3 expression in mouse pancreatitis specimens. Mice were treated with cerulein twice daily for 5 days to induce pancreatitis. There was increased expression of Par3 in mouse ADM lesions compared to the surrounding pancreas (**Fig. 1C**).

**Figure 1:**
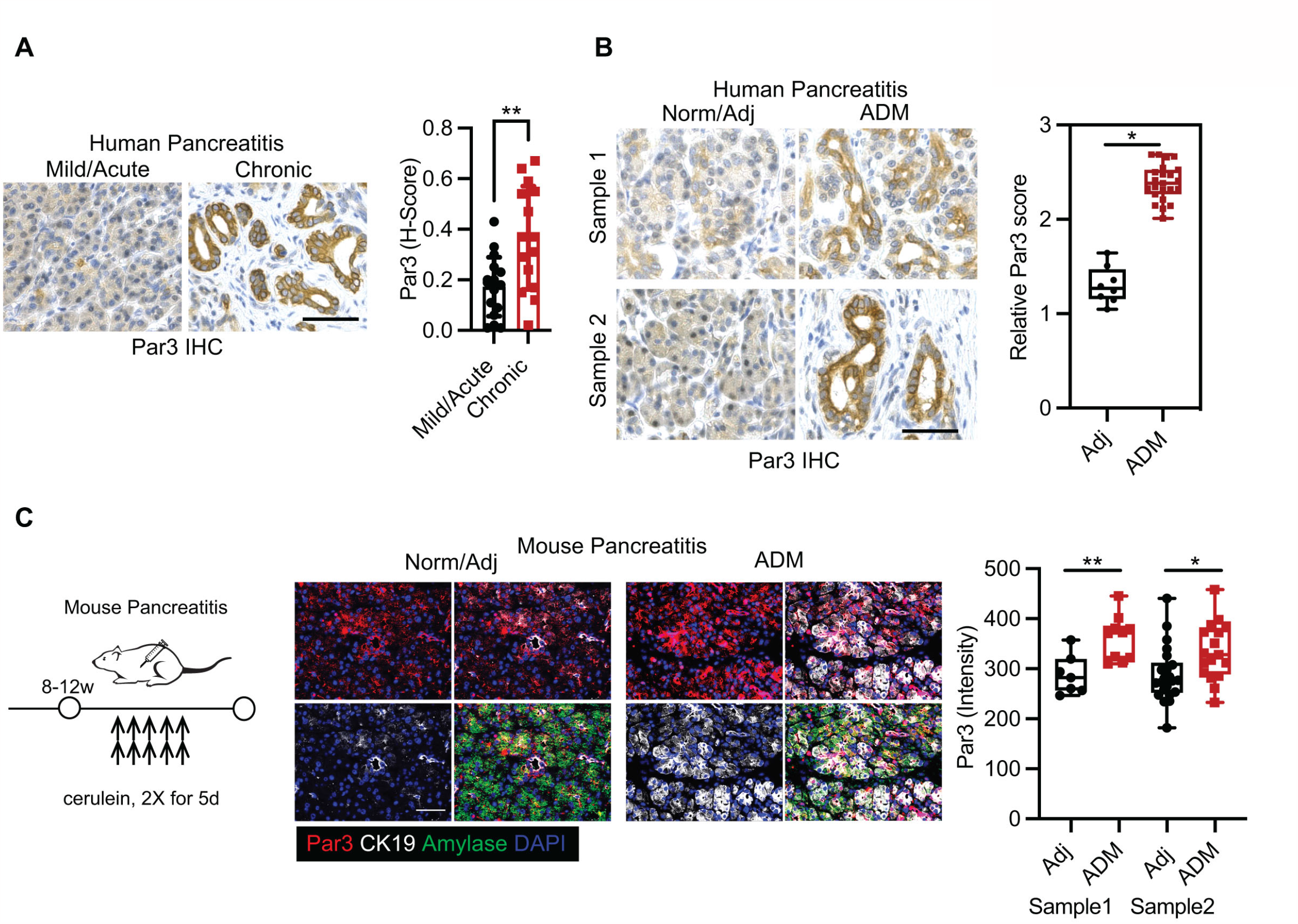
Increased Par3 expression in human and mouse pancreatitis specimens. (**A** and **B**) Human pancreatitis specimens were stained for Par3 by IHC. Par3 expression in the mild/acute pancreatitis was compared to chronic pancreatitis samples. The relative expression of Par3 in acinar-ductal metaplasia (ADM) compared to adjacent normal. (**C**) Wildtype B6 mice were treated with cerulein twice daily for 5 days to induce chronic pancreatitis, and the pancreas IF stained for Par3, DAPI, CK19, and amylase. Scale bar = 1 mm.

### Mice with pancreatic Par3 loss exhibit low-grade inflammation and acinar cell loss with aging

To evaluate the role of Par3 in vivo, we crossed Par3fl/+ mice with Pdx1-Cre mice to generate Pdx1-<u>C</u>re x Par3fl/+ (CPar3fl/+) mice, which were then further crossed to obtain Pdx1-<u>C</u>re x Par3fl/fl (CPar3fl/fl) mice (**Fig. 2A**). There was a decreased pancreatic expression of Par3 and the tight junction protein Zo-1 in the CPar3fl/fl mice (**Fig. 2B**). However, Par3 loss did not affect the expression of the adherens junction protein E-cadherin in the CPar3fl/fl mice (**Fig. 2B**). At three and six months of age, there was minimal to no difference in the body weights or the pancreas weights between the CPar3+/+ mice and the CPar3fl/fl mice (**Fig. 2C**). However, while there was not a gross difference in the acinar architecture, ductal structures, or pancreatic islets between the CPar3fl/fl and the CPar3+/+ mice at three months of age, there was evidence for loss of acinar cell mass in the CPar3fl/fl at six months of age (**Fig. 2D**). Moreover, at 12 months of age, more than 50% (5/9) of the CPar3fl/fl mice exhibited loss of acinar cells and increase in fat accumulation in the pancreatic parenchyma (pancreatic lipomatosis; **Fig. 2E**). When we evaluated for inflammation, there was an increased presence of macrophages in the CPar3fl/fl mice at three months of age, which continued to increase at six months of age (**Fig. 2F**).

**Figure 2:**
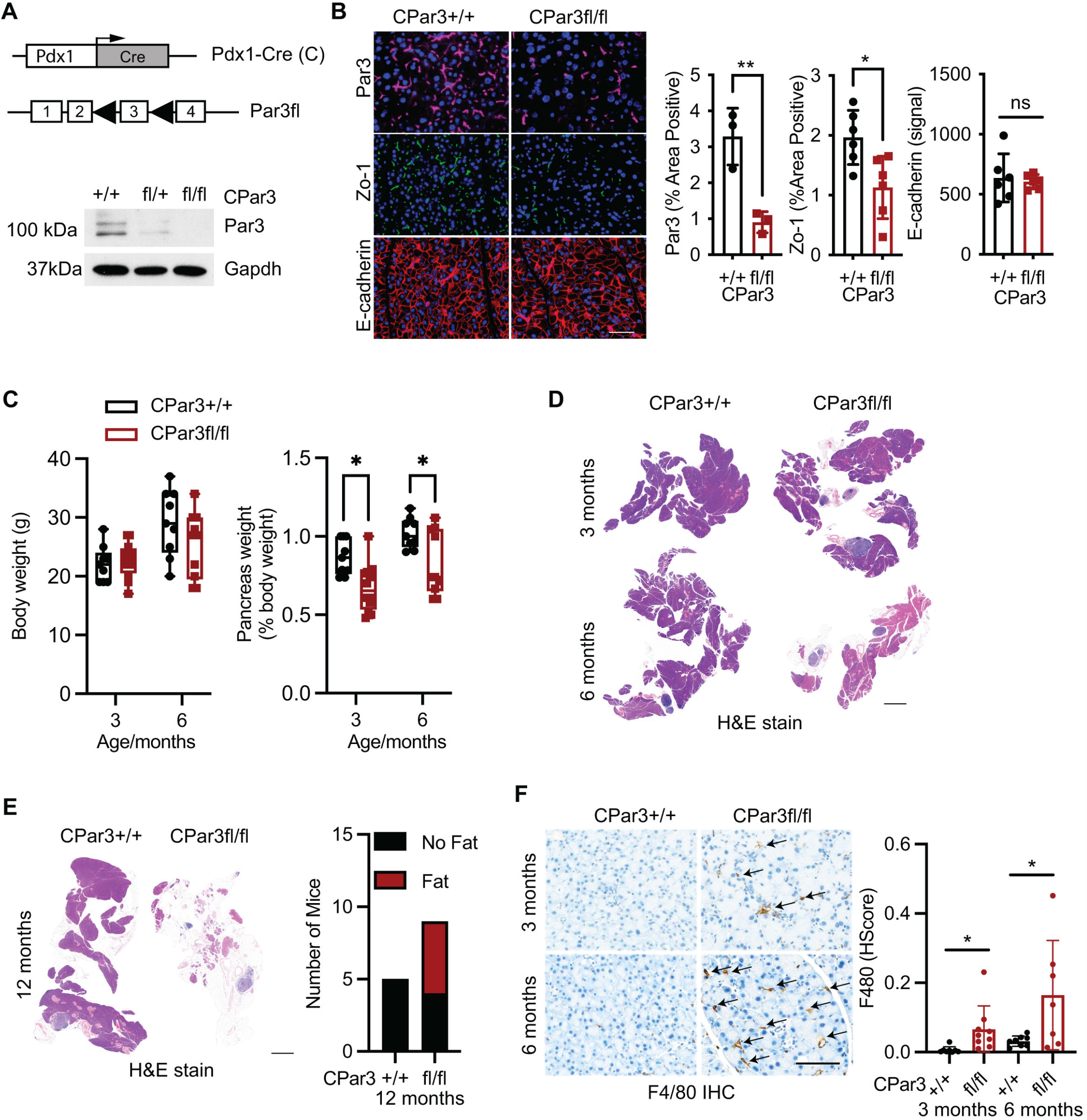
Mice with pancreatic Par3 loss exhibit low-grade inflammation and acinar cell loss with aging. (**A**) Alleles of Pdx1-Cre (C) and Par3fl mice. Par3 expression in the pancreas of mice of indicated genotypes was determined by Western blotting. (**B**) IF stains for Par3, Zo-1, and E-cadherin of the pancreas collected from CPar3+/+ and CPar3fl/fl mice at 13 weeks. Scale bar = 50 µm. Quantification of Par3 IF stains (n=3, 3). t-test, mean ±SD; ^*^ p-value ≤ 0.05. (**C**) Body weights and pancreas weights of CPar3+/+ and CPar3fl/fl mice at 3 and 6 months of age. (**D** and **E**) H&E stains of the pancreas from CPar3+/+ and CPar3fl/fl mice at 3, 6, and 12 months of age. Scale bar = 1 mm. (**F**) F4/80 stains of the pancreas from CPar3+/+ and CPar3fl/fl mice at 3 and 6 months of age. Scale bar = 100 µm.

### Pancreatic Par3 loss exacerbates cerulein-induced inflammation, acinar cell loss, and pancreatic lipomatosis

Given the worsening of the low-grade pancreatic inflammation and the development of lipomatosis in the CPar3fl/fl mice with aging, we evaluated whether exacerbating inflammation in young CPar3fl/fl mice would accelerate pancreatic lipomatosis. Thus, we evaluated the effects of Par3 loss on acinar cell injury with chronic pancreatitis in 2-3-month-old mice (**Fig. 3A**). While both CPar3+/+ and CPar3fl/fl mice lost weight with cerulein treatment, the CPar3fl/fl mice showed a more pronounced weight loss and loss of pancreas weight (**Fig. 3B**). When we examined the effects of Par3 loss on basal and cerulein-induced inflammation by staining for macrophages (F4/80) after five days of cerulein treatment, we found there were more macrophages in the pancreas of CPar3fl/fl mice than that of CPar3+/+ mice (**Fig. 3C**). H&E staining showed disruption of acinar parenchyma and loss of zymogen granules in the CPar3fl/fl mice after five days, loss of acinar cells after ten days, and a near-total absence of acinar cells after 14 days of cerulein treatment (**Fig. 3D**).

**Figure 3:**
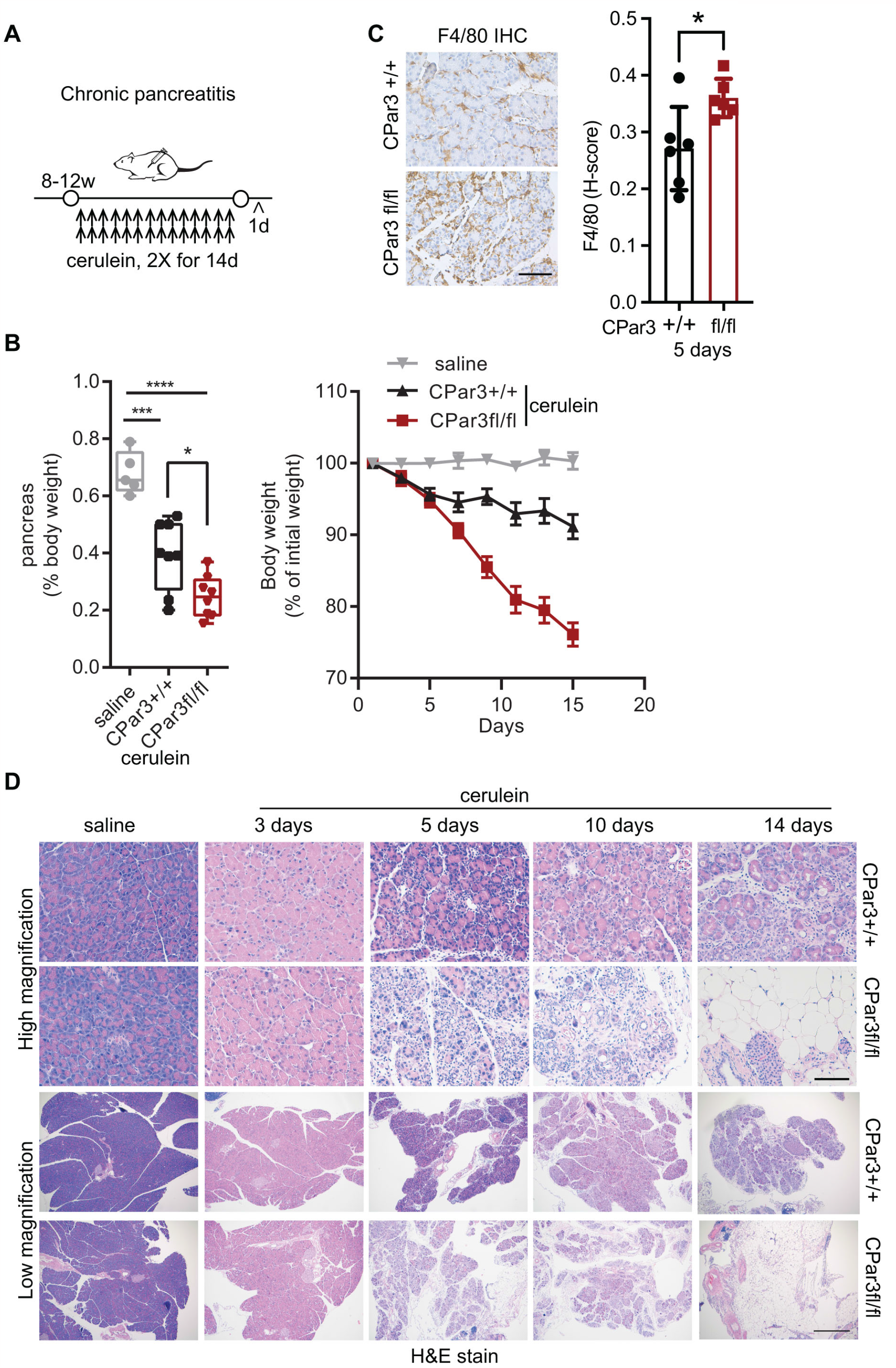
Pancreatic Par3 loss exacerbates cerulein-induced acinar cell loss and inflammation. (**A**) CPar3+/+ and CPar3fl/fl mice were treated with i.p. injections of saline or cerulein (250 µg/kg) twice daily for 14 days. (**B**) Weights of mice treated with saline (n=5) or cerulein (CPar3+/+, n=12; CPar3fl/fl, n=11) during treatment. One-way ANOVA, mean ±SD; ^**^ p-value ≤ 0.01, ^****^ p-value ≤ 0.0001. Pancreas weights as a percentage of body weight were determined for mice treated with saline (n=5) or cerulein (CPar3+/+, n=8; CPar3fl/fl, n=8). One-way ANOVA, mean ± min to max; ^*^ p-value ≤ 0.05, ^***^ p-value ≤ 0.001, ^****^ p-value ≤ 0.0001. (**C**) Pancreas from CPar3+/+ and CPar3fl/fl mice treated with cerulein for 5 days were stained for F4/80 (n=6,6), and the relative staining was quantified. t-test, mean ±SD; ^*^ p-value ≤ 0.05. Scale bar = 100 µm. (**D**) CPar3+/+ and CPar3fl/fl mice were treated twice daily for 3 (n=6,5), 5 (n=7,7), 10 (n=3,6), and 14 (n=4,5) days and the pancreas H&E stained. Scale bar = 100 µm.

Given the pronounced fatty replacement seen at Day 14 in the cerulein-treated CPar3fl/fl mice, we stained the Day 10 pancreatic specimens with Oil Red O to visualize fat. In contrast to the absence of Oil Red O staining in the pancreatic specimens from cerulein-treated CPar3+/+ mice, there was Oil Red O staining in pancreatic samples from cerulein-treated CPar3fl/fl mice (**Supplemental Fig. S1A**). After ten days of treatment, we also stained the pancreatic specimens for acinar (amylase) and ductal (CK19) markers. In both CPar3+/+ and CPar3fl/fl mice, the cerulein treatment increased the expression of CK19 (**Supplemental Fig. S1B**). However, compared to the cerulein-treated CPar3+/+ mice, there was a more pronounced reduction in amylase staining in the cerulein-treated CPar3fl/fl mice (**Supplemental Fig. S1B**), indicating extensive acinar cell loss. When we evaluated the effects of Par3 loss on apoptosis (cleaved-caspase 3, c-C3), there was increased c-C3 staining in the CPar3fl/fl mice following cerulein treatment (**Supplemental Fig. S1C**). These results suggest that Par3 restrains pancreatitis-induced acinar injury.

### Increased Par3 expression in mouse and human PanIN lesions

In parallel studies, we evaluated the role of Par3 in mutant Kras-induced phenotypic changes. Initially, we assessed the effects of mutant Kras on Par3 expression in mouse PanIN lesions ^17,18^. For these studies, we used LSL-<u>K</u>ras^**G12D**^/Pdx1-<u>C</u>re (KC) mice for early PanIN lesions and the KC/LSL-Tr<u>p</u>53^**R172H/+**^ (KPC) mice for more advanced PanIN lesions ^12,19^. Compared to control B6 mice, we found increased Par3 expression in the PanIN lesions present in the KC and the KPC mice (**Fig. 4A**). Moreover, there was increased expression of Par3 in mouse PanIN lesions compared to the surrounding pancreas in the KC mice (**Fig. 4B**). We also found increased Par3 expression in human PanIN lesions compared to the adjacent normal pancreas (**Fig. 4C**).

**Figure 4:**
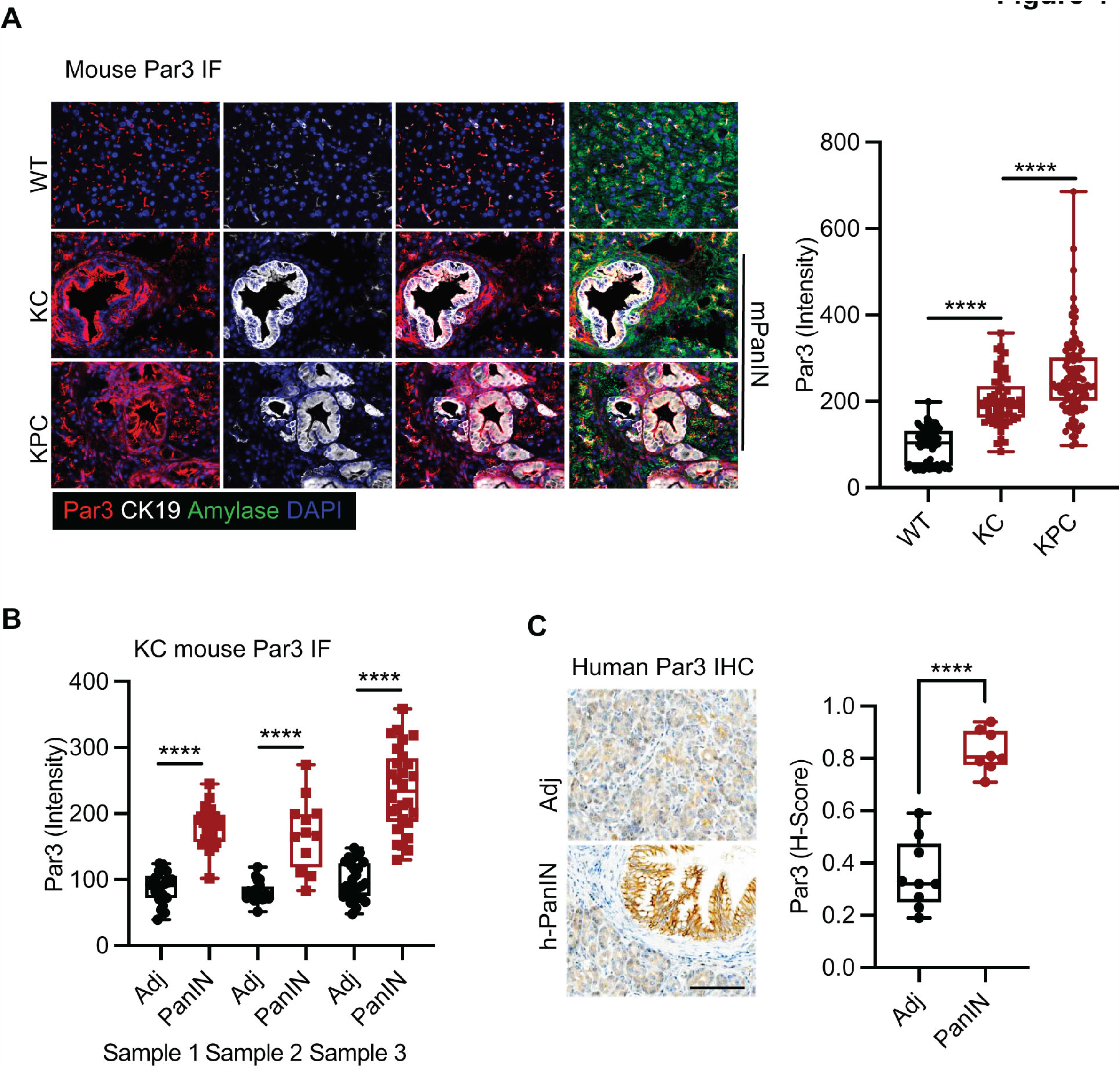
Increased Par3 expression in mouse and human PanIN lesions. (**A**) Pancreas sections from wild-type (WT) mice and PanIN lesions from KC (Kras) and KPC (Kras/p53) mice were IF stained for Par3, DAPI, CK19, and amylase and the relative Par3 expression in the PanIN lesions present in these mice was analyzed. (**B**) The Par3 expression in the PanIN lesions and the adjacent normal pancreas in the KC mice was quantified (**C**) Human pancreas specimens were IHC stained for Par3 and the relative Par3 expression in the PanIN lesions and the adjacent normal pancreas was quantified.

### Par3 loss in the KC model results in extensive PanIN lesions and large pancreatic cysts

To evaluate the role of Par3 in the KC mouse model, we crossed the LSL-<u>K</u>ras^**G12D**^ mice with the <u>C</u>Par3fl/fl mice to generate KCPar3fl/+ mice, which were then further crossed to obtain KCPar3fl/fl mice (**Fig. 5A**). Compared to the control KCPar3+/+ mice, the KCPar3fl/fl mice stopped gaining weight around 6-7 weeks of age (**Fig. 5B**). As demonstrated by Alcian blue staining, there was almost total involvement of the pancreas with PanIN lesions in the KCPar3fl/fl mice (**Figs. 5C** and **5D**). The KCPar3fl/fl mice developed pancreatic cysts, which, in some cases, were large enough to cause abdominal distension (**Figs. 5E** and **5F**). Consequently, the KCPar3fl/fl exhibited reduced survival compared to the control mice (**Fig. 5G**). These results suggest that Par3 restrains mutant Kras-driven phenotypic changes in the mouse pancreas.

**Figure 5:**
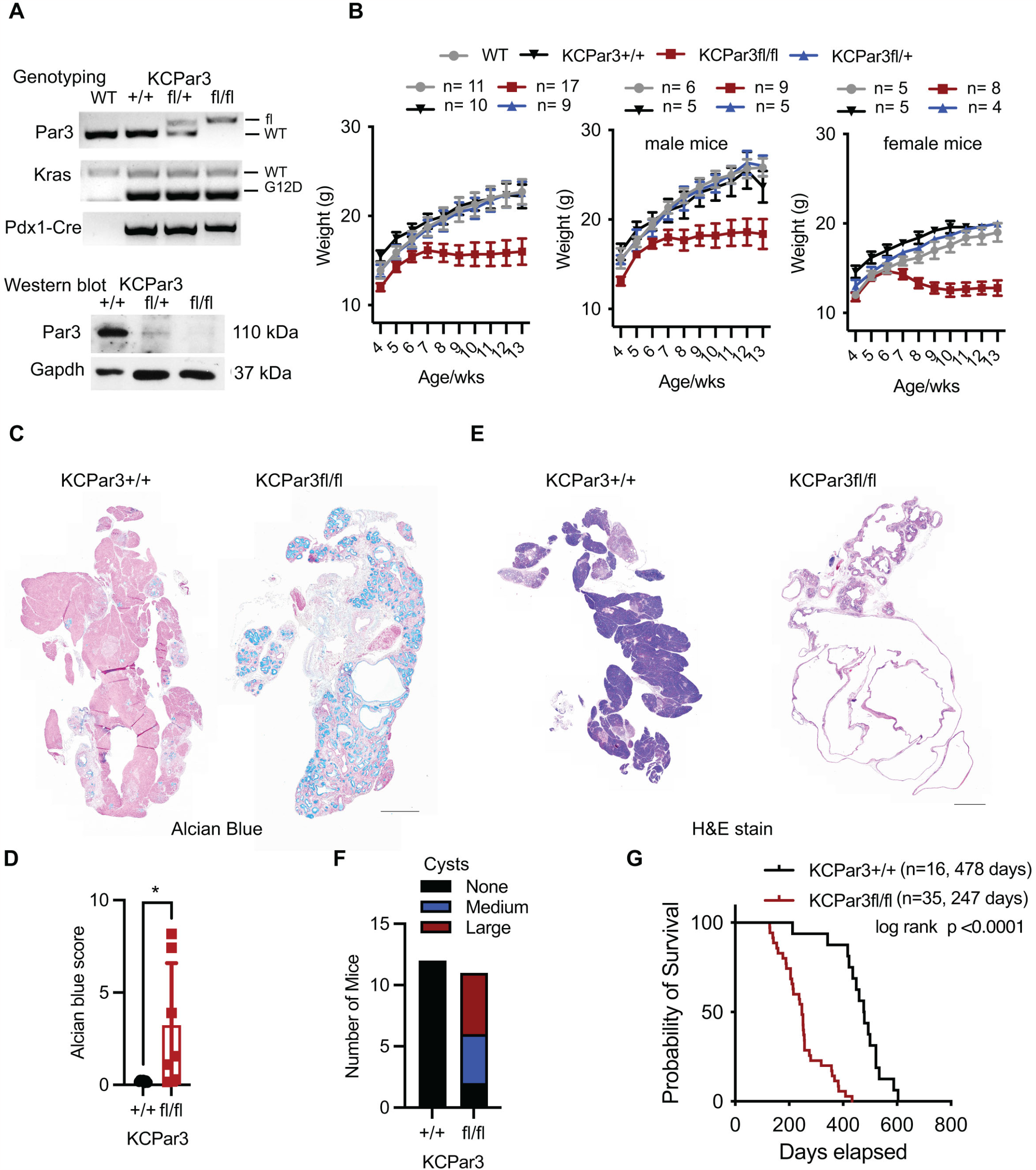
Par3 loss in the KC model results in extensive PanIN lesions and large pancreatic cysts. (**A**) Alleles of KrasG12D (K), Pdx1-Cre (C), and Par3fl mice. CPar3fl/+ mice were crossed with LSL-<u>K</u>ras (K) mice to create KCPar3fl/+ mice, which were then further crossed to generate KCPar3fl/fl mice. Genotyping PCR from tail DNA and Western blot for Par3 in the pancreas of these mice. (**B**) Weights of KCPar3+/+, and KCPar3fl/+, and KCPar3fl/fl mice were monitored till 13 weeks of age. (**C** and **D**) Alcian blue stain (to detect mucin present in PanINs) of the pancreas from 5-6-month-old KCPar3+/+ and KCPar3fl/fl mice. The relative staining of Alcian blue stain was quantified. (**E** and **F**) H&E stain of the pancreas from 6-month-old KCPar3+/+ and KCPar3fl/fl mice. Number of mice with no cysts or with medium and large pancreatic cysts. (**G**) Kaplan-Meier survival analysis of KCPar3+/+ (n=16) and KCPar3fl/fl (n=35) mice using log-rank test.

### Mice with pancreatic Par3 loss demonstrate reduced pancreatitis-induced primary cilia

To understand how Par3 loss exacerbates phenotypic changes in our mouse models, we performed RNA Sequencing and GSEA of the Day 5 pancreatitis samples. The GSEA showed a reduction in the ciliogenesis program in the CPar3fl/fl mice compared to the CPar3+/+ mice (**Figs. 6A** and **6B**). When we stained for primary cilia using acetylated tubulin and Arl13b, we found a significant reduction in the acetylated tubulin and Arl13b in the pancreas of cerulein-treated CPar3fl/fl mice compared to the pancreas of cerulein-treated CPar3+/+ mice (**Figs. 6C** and **6D**). Furthermore, when we stained the pancreas for FoxJ1, one of the key regulators of primary cilia ^20-22^, we found that cerulein treatment induced FoxJ1 in the CPar3+/+ mice but the cerulein-treated CPar3fl/fl mice had a near absence of FoxJ1 (**Fig. 6E**). Importantly, previous reports indicate that loss of pancreatic primary cilia results in pancreatitis, fatty changes, and cyst formation ^23,24^.

**Figure 6:**
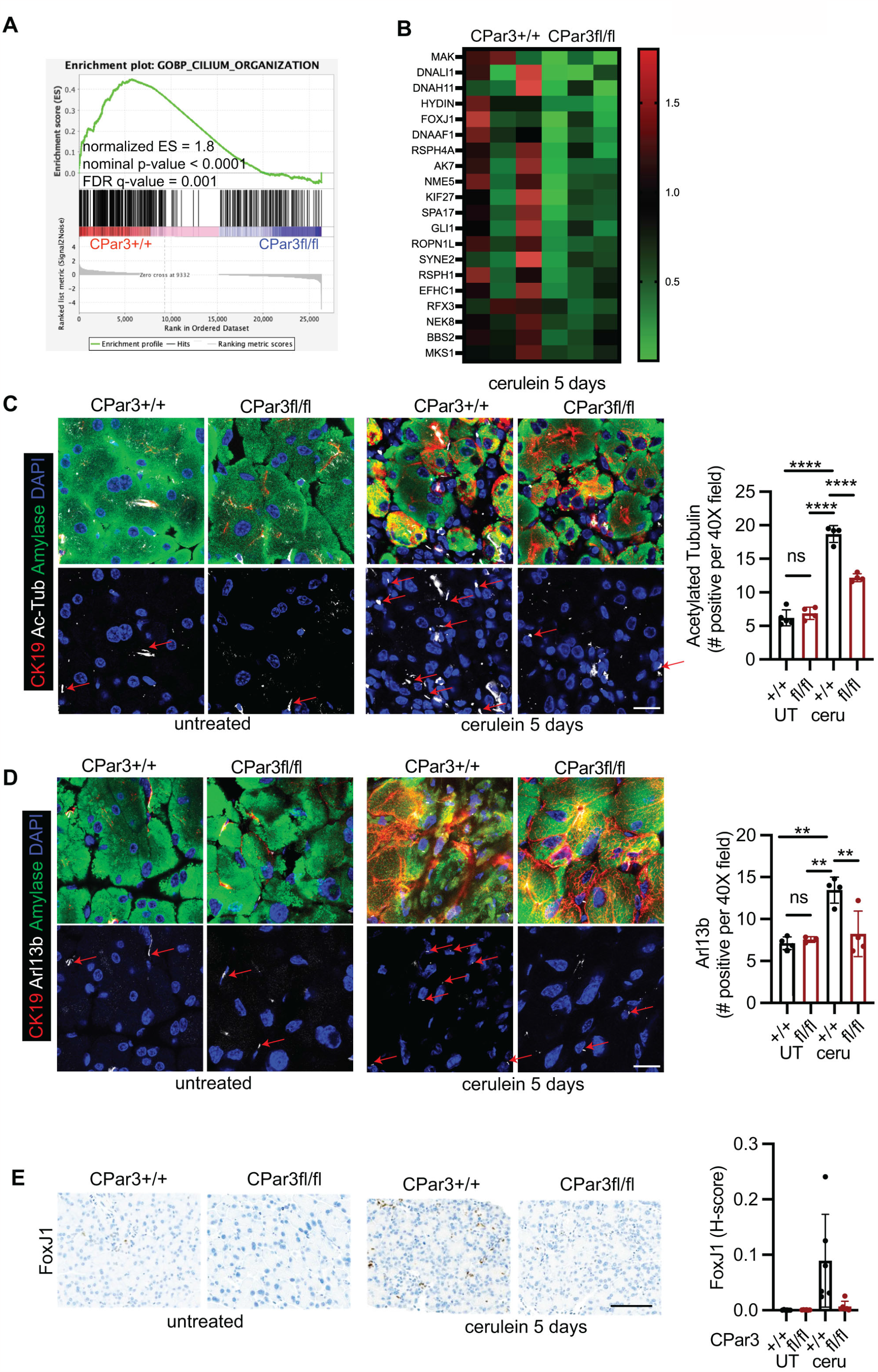
Mice with pancreatic Par3 loss demonstrate reduced pancreatitis-induced primary cilia. (**A** and **B**) CPar3+/+ and CPar3fl/fl mice were treated with i.p. injections of cerulein (250 µg/kg) twice daily for 5 days. Enrichment plot and GSEA for primary cilium using Day 5 pancreatitis RNA samples from CPar3+/+ and CPar3fl/fl mice (n=3,3). (**C-E**) Pancreas from control (UT) CPar3+/+ and CPar3fl/fl mice or from mice treated with cerulein for 5 days were stained for acetylated tubulin (n=5,4,4,4), Arl13b (n=4,3,4,4), and FoxJ1 (n=5,5,5,5), and the relative staining was quantified. Scale bar = 100 µm. t-test or One-way ANOVA, mean ±SD; ^**^ p-value ≤ 0.01, ^****^ p-value ≤ 0.0001.

### Targeting BET proteins attenuates pancreatitis-induced loss of primary cilia in mice lacking pancreatic Par3

As the BET protein Brd4 regulates acinar regeneration following pancreatitis-induced injury ^25^, we evaluated the effect of BET inhibition on pancreatitis-induced loss of primary cilia in the CPar3fl/fl mice. The CPar3fl/fl mice were pretreated with the BET inhibitor JQ1 for one week and then treated concurrently with JQ1 and cerulein for five days. Compared to the control (DMSO) cerulein-treated CPar3fl/fl mice, there was an increase in the ciliogenic program in the CPar3fl/fl mice co-treated with JQ1 and cerulein (**Fig. 7A**). While JQ1 did not attenuate cerulein-induced inflammation (**Fig. 7B**), JQ1 enhanced the expression of FoxJ1, acetylated tubulin, and Arl13b in the cerulein-treated CPar3fl/fl mice (**Figs. 7C-E**). These results indicate that BET inhibitors mitigate the effects of Par3 loss on pancreatitis-induced primary ciliogenesis program.

**Figure 7:**
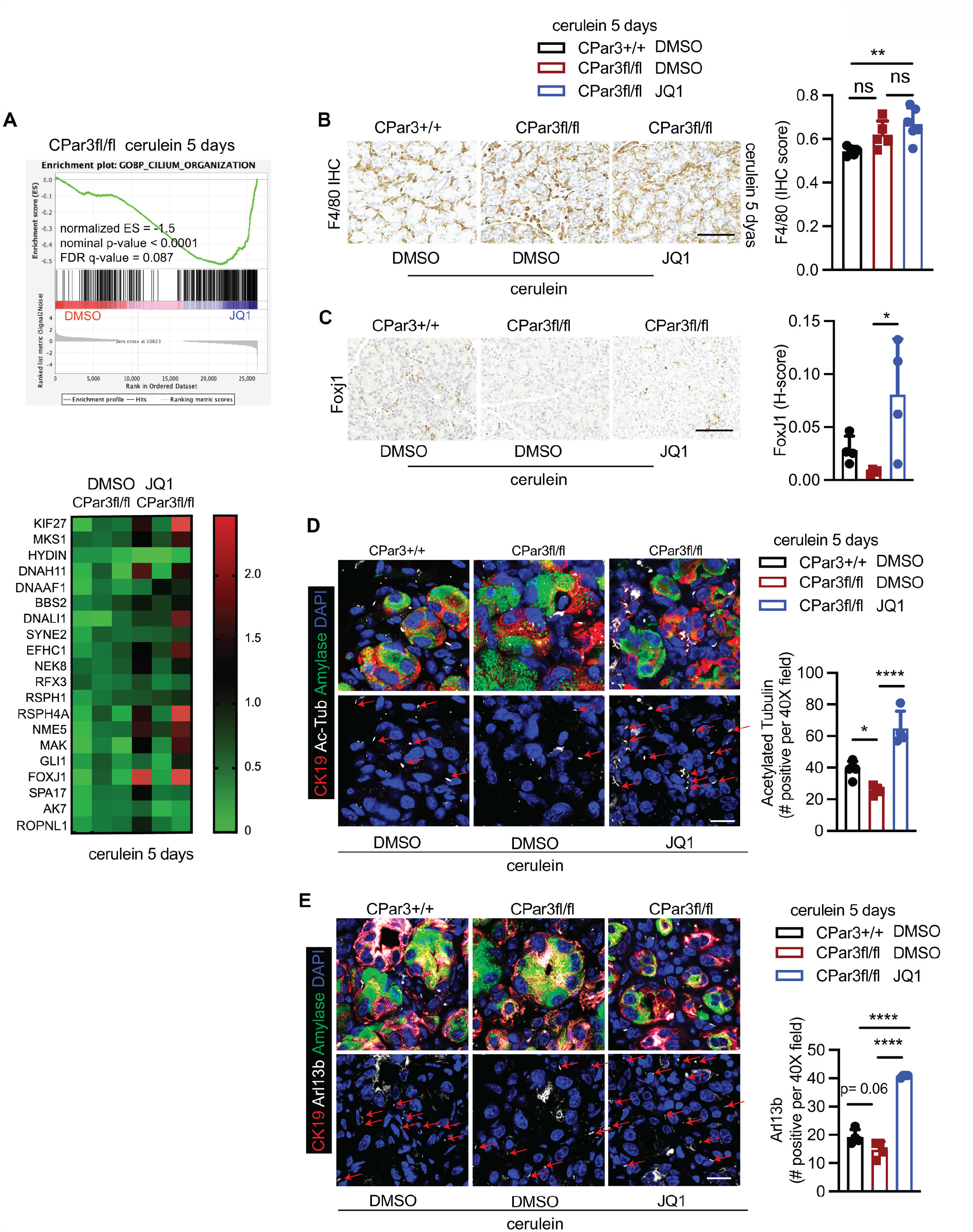
BET inhibitors attenuate pancreatitis-induced loss of primary cilia in mice lacking pancreatic Par3. CPar3+/+ and CPar3fl/fl mice were treated with DMSO or JQ1 (50mg/kg) for 7 days and then concurrently treated with DMSO or JQ1 (50 mg/kg) and cerulein (250 µg/kg) for additional 5 days. (**A**) Enrichment plot for primary cilium using RNA samples from CPar3fl/fl mice treated with cerulein and co-treated with DMSO or JQ1 (n=3,3). (**B**) Pancreas were stained for F4/80 and the relative staining was quantified (n=5,5,6). (**C-E**) Pancreatic section from mice treated with cerulein for 5 days and co-treated with DMSO or JQ1 were stained for FoxJ1 (n=4,4,5), acetylated tubulin (n=5,4,4), and Arl13b (n=4,4,4) and the relative staining was quantified. Scale bar = 100 µm. One-way ANOVA, mean ±SD; ^*^ p-value ≤ 0.05, ^**^ p-value ≤ 0.01.

### Targeting BET proteins attenuates cerulein-induced acinar cell loss in mice lacking pancreatic Par3 and enhance recovery of acinar cell mass and body weight

Initially, we evaluated whether the cerulein-treated CPar3fl/fl mice could recover normal pancreas architecture after stopping cerulein treatment. While the control CPar3+/+ mice quickly regained weight after stopping cerulein treatment, the CPar3fl/fl mice regained weight at a slower rate and failed to fully recover the lost weight 14 days after stopping cerulein treatment (**Supplemental Fig. S2**). There was also minimal to no recovery of the acinar compartment in the CPar3fl/fl mice after stopping cerulein treatment (**Supplemental Fig. S2**).

Finally, we evaluated the effects of BET inhibitors on cerulein-induced acinar cell loss in the CPar3fl/fl and the CPar3+/+ mice. JQ1 did not prevent weight loss in cerulein-treated CPar3+/+ and CPar3fl/fl mice (**Fig. 8A**). However, histologic examination showed preservation of the acinar compartment in the CPar3fl/fl mice, as demonstrated by amylase staining (**Fig. 8B**). Consequently, pre-treatment with JQ1 accelerated the recovery of body mass of both CPar3+/+ and CPar3fl/fl mice (**Fig. 8C**). Histologic examination showed significant recovery of the acinar compartment of CPar3fl/fl mice, as demonstrated by amylase staining (**Fig. 8D**).

**Figure 8:**
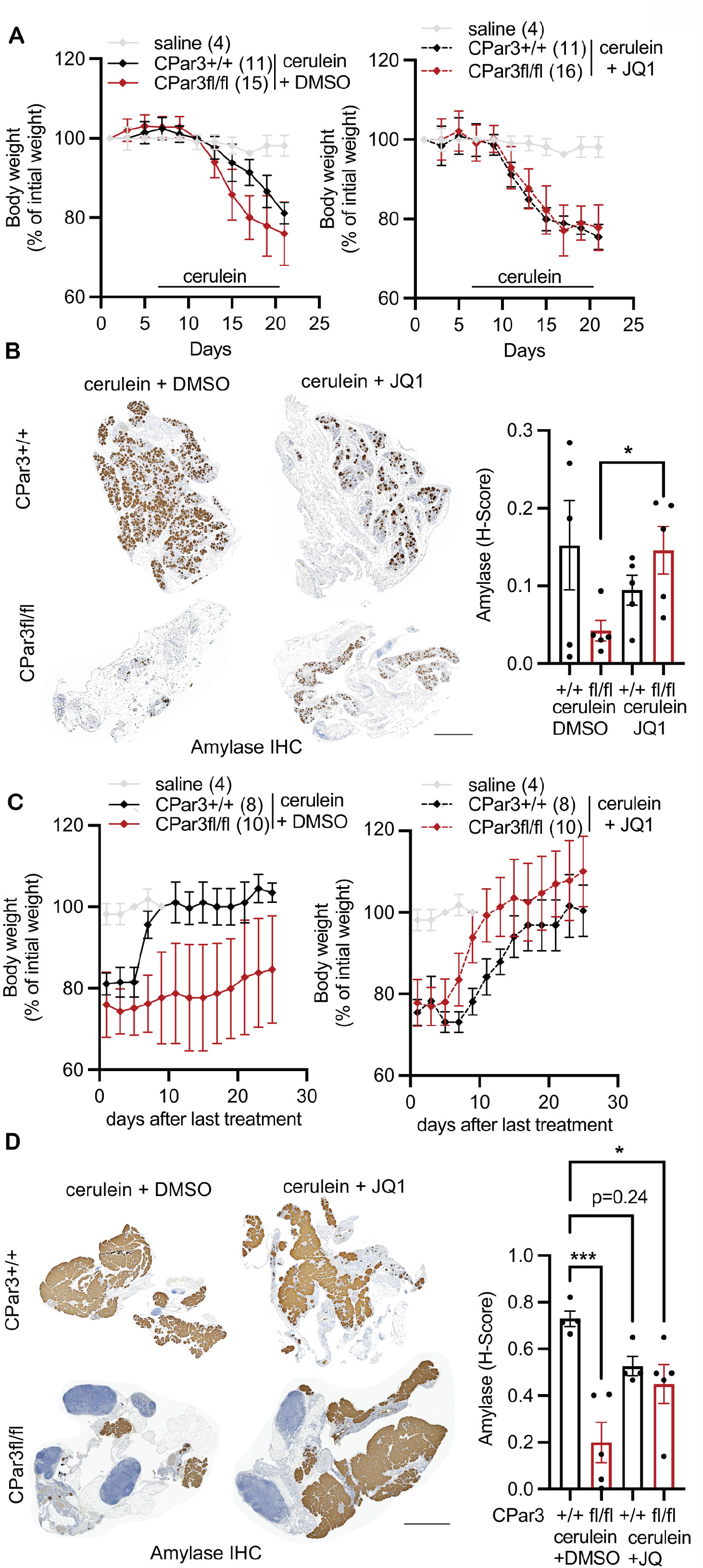
BET inhibitors attenuate cerulein-induced acinar cell loss in mice lacking pancreatic Par3 and enhance recovery of acinar cell mass and body weight. (**A** and **B**) CPar3+/+ and CPar3fl/fl mice were treated with DMSO or JQ1 (50mg/kg) for 7 days and then concurrently treated with DMSO or JQ1 (50 mg/kg) and cerulein (250 µg/kg) for additional two weeks. Mice were monitored for weight loss during treatment. Acinar cell loss at the end of treatment was analyzed by amylase staining (n=5 for CPar3+/+ and CPar3fl/fl samples), and the relative amylase staining was quantified. One-way ANOVA, mean ±SD. ^*^ p-value ≤ 0.05. Scale bar = 1 mm. (**C** and **D**) CPar3+/+ and CPar3fl/fl were pre-treated with DMSO or JQ1 (50mg/kg) for 7 days and then concurrently treated with DMSO or JQ1 (50mg/kg) and cerulein (250 µg/kg) for two weeks. Mice were then monitored for additional 25 days. Recovery of body weights monitored over 25 days. Acinar cell recovery 25 days post-treatment was analyzed by amylase staining (n=4 for CPar3+/+ and n=5 for CPar3fl/fl samples), and the relative amylase staining was quantified. One-way ANOVA, mean ±SD; ^*^ p-value ≤ 0.05, ^***^ p-value ≤ 0.001. Scale bar = 1mm.

## DISCUSSION

The polarity proteins have been studied extensively in tumorigenesis ^26,27^; however, their role in normal tissue homeostasis has yet to be fully understood. For example, loss of Par3 in the prostate tissue promotes proliferation and causes high-grade prostatic intraepithelial neoplastic lesions ^28^. Similarly, the loss of Par3 in the epidermis causes aberrant proliferation and differentiation ^6^. Moreover, targeting Par3 in mouse mammary progenitor cells causes ductal hyperplasia and disorganized end bud structures that fail to remodel into normal ductal structures when transplanted ^29^. In contrast to these studies, we show that loss of Par3 in the pancreas does not affect pancreas development.

However, one of the most dramatic phenotypes in mice with pancreatic Par3 loss is the extensive loss of the acinar compartment with aging and following induction of pancreatitis. We show there is increased apoptosis following pancreatitis-induced tissue injury. Loss of epidermal Par3 is also associated with increased apoptosis following treatment with DMBA/TPA ^30^. Par3 in the epidermis maintains mitotic accuracy and cellular fitness by integrating biomechanical signaling and cell division ^7^. These results suggest that the disruption of apical-basal polarity induces apoptosis, possibly as a protective mechanism against transformation in damaged tissue. We also show that in mice with pancreatic Par3 loss, there is increased inflammation following the induction of pancreatitis and with aging. Notably, mice lacking Par3 in the endothelium demonstrate increased macrophage infiltration in the aortic arch ^31^, indicating that Par3 limits endothelial inflammation in blood vessels under laminar flow ^31^. These results suggest that Par3 restrains tissue injury and inflammation in multiple organ systems to maintain tissue homeostasis.

We found induction of pancreatitis in mice with Par3 loss results in the loss of primary cilia. Our findings agree with a recent study showing that loss of Par3 in neural precursors *in vivo* perturbs the integrity of primary cilia ^32^. Also, an earlier study showed that Par3 is required for the development of primary cilia in MDCK cells ^33^. Notably, a lack of primary cilia in the pancreas results in severe pancreatic abnormalities, including acinar cell loss and associated lipomatosis ^23,24^. Lack of primary cilia is also associated with cyst formation in the pancreas ^23,34,35^, as well as in the kidney and liver ^36,37^. We have found that Par3 loss also results in cyst formation in the mutant Kras-driven mouse model of pancreatic tumor initiation. In contrast to the loss of primary cilia in human PanIN lesions ^38^, we have found that Par3 expression is increased in mouse and human PanIN lesions. However, the loss of Par3 in the mutant Kras-driven mouse model results in extensive PanIN formation, suggesting that Par3 restrains mutant Kras-driven phenotypic changes in the mouse pancreas.

We show that mice with Par3 loss demonstrate minimal to no recovery of the acinar compartment following induction of pancreatitis. However, targeting BET proteins attenuates cerulein-induced acinar cell loss and enhances acinar cell mass recovery in these mice. The Scott Lowe lab had previously shown that genetic targeting of Brd4 using shRNA enhanced pancreatitis-induced acinar injury but prevented subsequent regeneration ^25^. We have also found that targeting BET proteins with the BET inhibitor JQ1 exacerbated pancreatitis-induced injury in control mice. However, in mice with Par3 loss, while JQ1 did not attenuate pancreatitis-induced inflammation, JQ1 attenuated pancreatitis-induced acinar cell loss. Also, when we stopped JQ1 treatment, in contrast to the persistent knockdown of Brd4 with shRNA ^25^, we found that both the control mice and mice with pancreatic Par3 loss demonstrated recovery of acinar cell mass and body weight. Unexpectedly, we found that JQ1 rescued chronic pancreatitis-induced primary cilia in mice with Par3 loss. Interestingly, several small molecule inhibitors induce ciliogenesis by modulating Aurora-A (AURKA) activity ^39,40^, one of the central regulators of ciliary disassembly ^41,42^. In future studies, we will evaluate how BET inhibitors promote primary cilia.

Overall, our study increases our understanding of the role of the polarity protein Par3 in restraining pancreatitis and mutant Kras-driven phenotypic changes and identifies BET inhibitors to attenuate pancreas injury and facilitate pancreas tissue regeneration.

## METHODS

### Animal experiments

#### Conditional knockout

Mice with loss of Par3 in the pancreas were generated by crossing Pdx1-Cre mice (Jackson Laboratory #014647) to mice expressing the floxed allele of *Par3* ^43,44^, to generate C•;Par3fl/+ and C•;Par3fl/fl mice. All mice were bred on a C57/BL6 background, and both genders were used in the studies.

#### Induction of pancreatitis

Pancreatitis was induced in age- and litter-matched 8-to 12-week-old mice of both genders. Chronic pancreatitis was induced by intraperitoneal injections of cerulein (250 µg/kg) two times daily for 3, 5, 10, and 14 days, and the pancreas was collected 16-20 hours later.

### Histology/Immunohistochemistry (IHC)

For immunostains, paraffin-embedded sections were deparaffinized and rehydrated. Antigen retrieval was performed by boiling for ten minutes in either Tris EDTA (10mM Tris 1mM EDTA, 0.05% Tween 20, pH 9.0) or sodium citrate buffer (10 mM sodium citrate, 0.05% Tween 20, pH 6.0) using a pressure cooker. Endogenous peroxidase activity in tissue was quenched, and sections were blocked with a mixture of goat serum and bovine serum albumin (BSA). Tissue sections were incubated with primary antibodies: F4/80 (Cell Signaling #70076, 1:800), cleaved caspase-3 (Cell Signaling Cat# 9664, 1:1000), Amylase (Cell Signaling #3796, 1:2000), Par3 (Millipore-Sigma #HPA030443, 1:400), and FoxJ1 (Abcam #ab235445, 1:500) overnight at 4°C. Antibody binding was detected using HRP-conjugated anti-rabbit secondary antibody (Vector Laboratories MP-7451) and visualized using ImmPACT DAB Peroxidase Substrate kit (Vector Laboratories, SK-4105). Photographs were taken on the TissueGnostics system and analyzed by ImageJ or Aperio Software (Leica Biosystems). Hematoxylin, and eosin (H&E) Oil RedO, and Alcian blue stains were performed by the Northwestern University Pathology Core Facility.

### Immunofluorescence

In select experiments, freshly isolated tissue samples were snap-frozen in OCT, and then sections were fixed in cold acetone:methanol (1:1) for 20 minutes. After antigen retrieval (similar to IHC), tissue sections were stained using the mouse-on-mouse (MOM) immunodetection kit (Vector Laboratories (BMK-2202). Tissue sections were blocked with MOM-blocking reagent for 1 hour at room temperature, followed by washing and incubation with MOM-diluent for 5 minutes. Tissue sections were then incubated with primary antibodies: Par3 (Millipore Sigma #07-330, 1:100), E-cadherin (BD Biosciences #610181, 1:500), Zo-1 (Santa Cruz Biotech #sc-33725, 1:200), Amylase (Santa Cruz Biotech #sc-46657, 1:500), CK19 (Developmental Studies Hybridoma Bank (DSHB) TROMAIII, 1:100), acetylated tubulin (acetyl K40) (Abcam #ab179484, 1:1000), and Arl13b (DSHB N295B/66 1:10) for 1 hour at room temperature or at 4 °C overnight. After washing, the sections were incubated with Alexa Fluor conjugated secondary antibodies (Thermo Fisher goat anti-rabbit AF488 Plus #A32731, goat anti-mouse AF647 #A21235, and goat anti-rat AF546 #A11081) for 30 minutes in the dark at room temperature. The tissue sections were washed and incubated with DAPI (Thermo Fisher D1306) for 10 minutes. Slides were mounted with fluorescence mounting media and images acquired using the EVOS M5000 epifluorescence microscope (Thermo Fisher). Images were also acquired using the Nikon A1 confocal laser scanning microscope equipped with four laser lines (405, 488, 561) and high sensitivity GaAsP detectors.

### Western blot

Tissue samples were finely ground and lysed with cold RIPA lysis buffer containing protease and phosphatase inhibitors. The lysates were then clarified by centrifugation at 10,000 rpm for 10 minutes at 4°C, and the protein concentration was determined using Precision Red solution (Cytoskeleton, Inc., Denver, CO). Equal amounts of protein were separated with a 10% SDS-PAGE electrophoresis gel. The separated proteins were transferred to a nitrocellulose membrane using the semi-dry transfer system (Bio-Rad). After blocking for one hour at room temperature with 5% BSA, the membranes were incubated overnight at 4°C with primary antibodies. Primary antibodies used include Par3 (Millipore Sigma #07-330, 1:1000) and Gapdh (Millipore Sigma #MAB374, 1:5000). HRP-conjugated rabbit (A6667) or mouse (A4416) secondary antibody (Millipore-Sigma St. Louis, MO) was used with SuperSignal West Pico PLUS (Thermo Fisher Scientific) for protein detection.

### RNAseq

RNA was extracted from mouse pancreas tissue using the RNeasy kit. The RNA was submitted to Active Motif, Inc (Carlsbad, California) for cDNA library preparation and sequencing. The FASTQ and BAM files along with analyses including differential analysis (DESeq2 software) and GSEA (MSigDB C5 GO gene set) results were provided. The raw FASTQ and processed files were deposited in the Gene Expression Omnibus (GEO) with accession number GSE243623.

### PCR

Genomic DNA was isolated from mouse tail biopsies using the GenElute Mammalian Genomic DNA extraction kit (Millipore-Sigma, G1N350-1KT) according to the manufacturer’s instructions. The REDTaq ReadyMix PCR reaction mix was use to amplify specific DNA fragments using primers for mutant Kras-G12D (wildtype: TGT CTT TCC CCA GCA CAG T, mutant: GCA GGT CGA GGG ACC TAA TA, and common: CTG CAT AGT ACG CTA TAC CCT GT), Pdx1-Cre (forward: CCT GGA AAA TGC TTC TGT CCG and reverse: CAG GGT GTT ATA AGC AAT CCC), Par3fl (forward: AGG CTA GCC TGG GTG ATT TGA GAC C and reverse: TTC CCT GAG GCC TGA CAC TCC AGT C), and p53-R172H (wildtype: TTA CAC ATC CAG CCT CTG TGG, mutant: AGC TAG CCA CCA TGG CTT GAG TAA GTC T, and common: CTT GGA GAC ATA GCC ACA CTG). The amplified DNA was resolved on an agarose gel and imaged using the ChemiDOC system (Bio-Rad).

### Statistics

Statistical details of experiments can be found in the figure legends, including the statistical tests used and the number (n) of animals. All statistical analyses were done using GraphPad Prism. The *in vivo* and *in vitro* results were compared using one-way ANOVA with Tukey correction or 2-tailed *t*-test analysis. Error bars represent standard deviation as specified in the figure legends. A p-value of less than 0.05 was considered significant.

### Study approval

The Northwestern University Institutional Animal Care and Use Committee approved all animal work and procedures. The animal experiments were performed per relevant guidelines and regulations. The animals were housed at 12h light/dark cycle in ventilated cages with controlled temperature and humidity. The animals were provided water and a standard mouse diet *ad libitum*, with bedding changed regularly.

## AUTHOR CONTRIBUTIONS

M.A.S. designed the studies, performed the experiments, analyzed the data, and wrote the manuscript. A.E.M., C.S., and T.N.D.P. performed the experiments and analyzed the data. T.H. and S.O. generously provided the Par3fl/fl mice. H.G.M. designed the studies, analyzed the data, wrote the manuscript, and secured funding. All authors edited and approved the final manuscript.

## FUNDING

This work was supported by grants R01CA217907 (to H.G.M.), R01CA265997 (to H.G.M.), a Merit award I01BX005595 (to H.G.M.) from the Department of Veterans Affairs, and APA/APA Foundation 2020 Young Investigator Pancreatitis Grant (to M.A.S.). The contents of this article are the responsibility of the authors and do not represent the views of the Department of Veterans Affairs or the United States Government.

## ACKNOWLEDGEMENTS

This work was supported by the Northwestern University Pathology Core Facility and the Northwestern University Center for Advanced Microscopy, both generously supported by NCI CCSG P30CA060553, awarded to the Robert H Lurie Comprehensive Cancer Center. We would like to thank Nida Mubin and Yaning Xi for their assistance with some of the experiments.

## SUPPLEMENTAL FIGURE LEGENDS

**Figure S1: Pancreatitis exacerbates loss of acinar compartment in mice with pancreatic Par3 loss**. (**A**) Pancreas from CPar3+/+ (n=3) and CPar3fl/fl (n=4) mice treated with cerulein for 10 days were stained for Oil Red O. Scale bar = 100 µm. (**B**) Pancreas from CPar3+/+ (n=3) and CPar3fl/fl (n=6) mice treated with cerulein twice daily for 10 days were stained for CK19 and amylase, and the relative staining was quantified. t-test, mean ±SD; ^*^ p-value ≤ 0.05, ^****^ p-value ≤ 0.0001. Scale bar = 50 µm. (**C**) Pancreas from CPar3+/+ and CPar3fl/fl mice treated with cerulein for 5 days were stained for cleaved caspase-3 (c-C3) (n=6,7), and the relative staining was quantified. t-test, mean ±SD; ^*^ p-value ≤ 0.05. Scale bar = 100 µm.

**Figure S2: Mice with pancreatic Par3 loss do not recover acinar cell mass and body weight following induction of chronic pancreatitis**. (**A** and **B**) CPar3+/+ and CPar3fl/fl mice were treated with saline or cerulein (250 µg/kg) twice daily for 14 days. Weights of mice treated with saline (n=4) or cerulein (CPar3+/+, n=7; CPar3fl/fl, n=9) during and after treatment. t-test, mean ±SD; ^*^ p-value ≤ 0.05. Amylase staining of the pancreas 2 weeks (saline, n=3; cerulein, CPar3+/+, n=4; cerulein, CPar3fl/fl, n=3) and 2 months (cerulein, CPar3fl/fl, n=5) post-cerulein treatment, and the relative amylase staining was quantified. One-way ANOVA, mean ±SD; ns, not significant, ^*^ p-value ≤ 0.05, ^**^ p-value ≤ 0.01. Scale bar = 1 mm.

## Notes

**Conflict of interest statement:** The authors have declared that no conflict of interest exists.

### Competing Interest Statement

The authors have declared no competing interest.

### Summary of Updates

Most of the data in Figure 6 in previous version has been changed to Figure 7. A new figure 6 includes data showing how loss of Par3 impacts ciliogenesis in pancreatitis. The updated manuscript now has 8 figures.

