## Supplementary figures and images for "BET inhibition rescues ciliogenesis and ameliorates pancreatitis-driven phenotypic changes in mice with Par3 loss" (from another version of this preprint: version 1, 2023-09-17)

### Supplementary figures

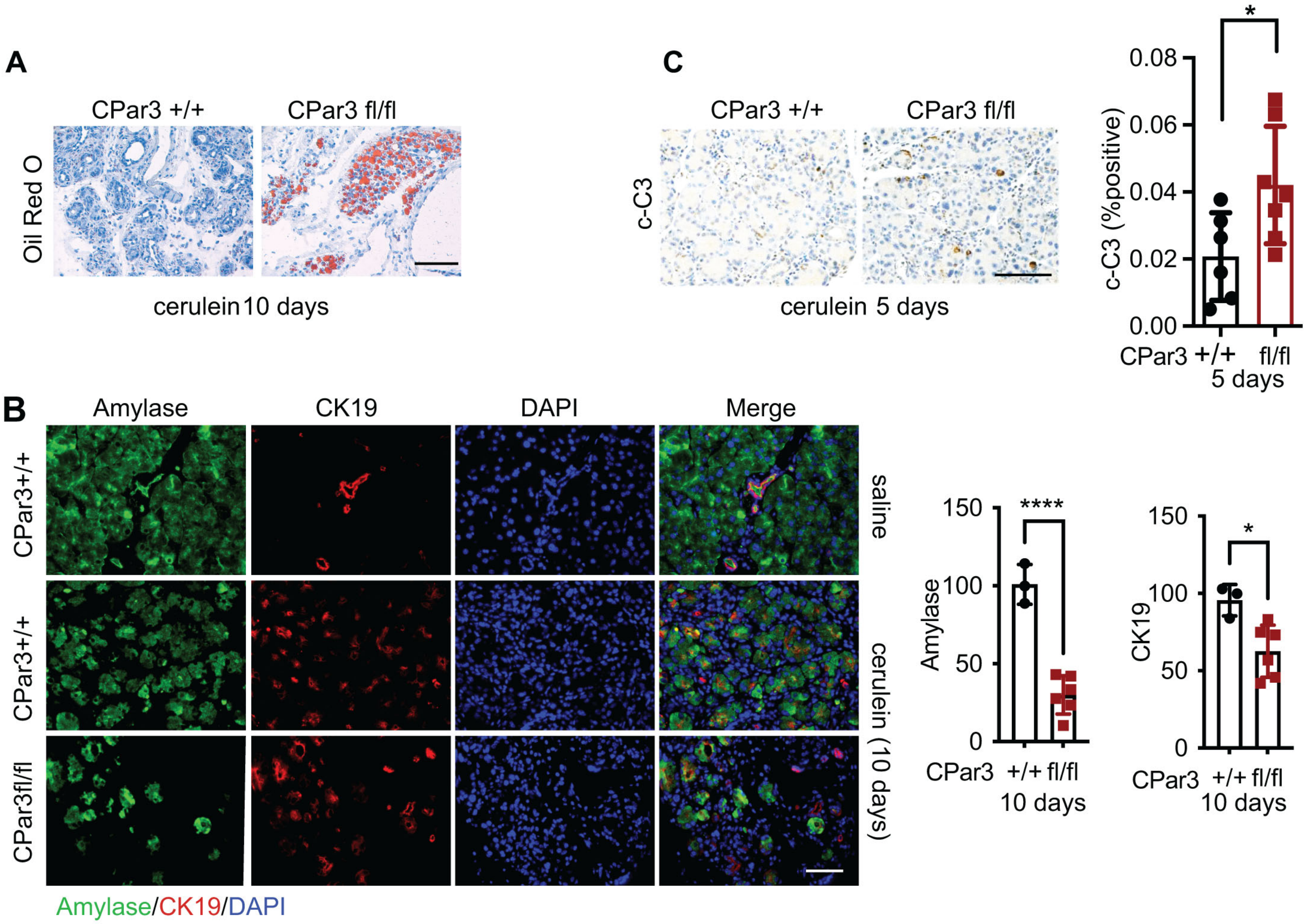

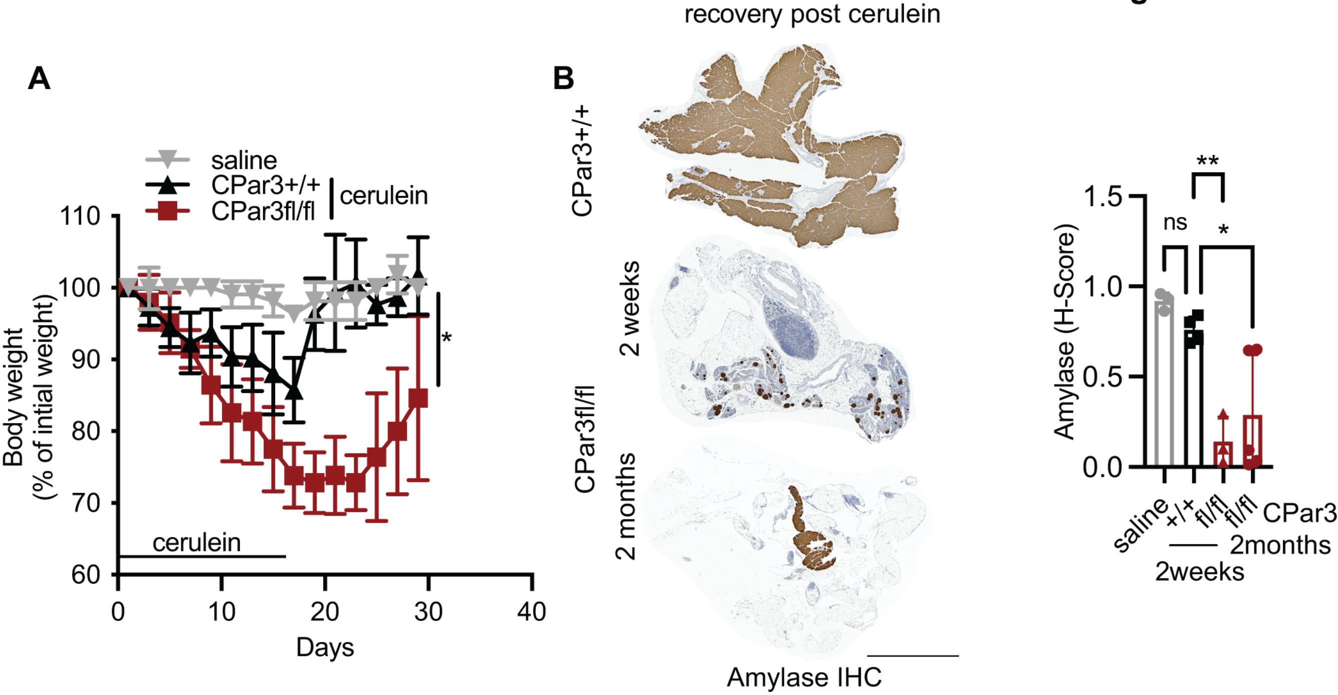
